# Close Packing of Cells in Vestibular Epithelia Supports Local Electrical Potentials that Reduce Latency of Action Potential Generation

**DOI:** 10.1101/2025.06.14.659580

**Authors:** Aravind Chenrayan Govindaraju, Robert M. Raphael

## Abstract

In the vestibular system, upon transduction of head motion, ionic currents from type I sensory hair cells alter [K^+^] and electrical potentials in an extended synaptic cleft formed by a calyx terminal of the associated afferent neuron. During excitatory stimuli, these changes in turn modulate post-synaptic currents across the calyx inner face to depolarize the afferent and initiate action potentials. Within the tightly packed columnar vestibular sensory epithelium, electrical currents from the hair cell and calyx must also traverse non-synaptic extracellular spaces and generate local extracellular potentials before dispersing into the perilymph beneath the basement membrane. Here we show that such dynamic electrical potentials enhance action potential generation by reducing outward K^+^ currents on both the inner and outer faces of the calyx. This effect also influences adjacent calyces and may explain the abundance of calyx terminals in amniotes where there is a need for rapid recognition of changes in head orientation and acceleration.

## Introduction

In amniote vestibular sensory epithelia, up to 74% of hair cells can be type-I (1), and upon transduction of head motion, transmit to a cup-like (*calyx*) terminal of an afferent neuron. Transmission to the afferent neuron can occur without neurotransmitter release (*non-quantal transmission, NQT*) (2). During NQT the modulation of potassium ion concentration ‘[K^+^]’ and electrical potential ‘ϕ’ in the narrow (3) synaptic cleft alter ionic currents on the post-synaptic terminal of the afferent neuron (4, 5). This process is facilitated by voltage-activated K^+^ channels present on the type-I hair cell, calyx terminal and afferent fiber (6–10) that typically mediate an efflux of K^+^ ions. K^+^ currents primarily flow out of the hair cell through a low voltage-activated potassium conductance comprised of K_V_1.8 subunits (11) and into the cleft through K_V_7 channels on the inner face of the calyx (3, 6, 7). Until now, the fast encoding of head motion by vestibular primary afferent neurons has mostly been attributed to processes and morphological features upstream from the afferent: variations in hair bundle and otolith morphology (12), rapid transmission across the synaptic cleft due to large hair cell currents (13), and the effect of synaptic K^+^ accumulation (10) on currents across the calyx inner face (*CIF*). However, while synaptic transmission can initiate afferent depolarization, AP generation is a function of ion channels on the afferent neuron. Calyx afferents can exhibit spontaneous firing independent of synaptic transmission, and AP firing rates change when K_V_1 and K_V_7 channels are blocked (8, 9, 14). Efferent activity also mediates slow excitation of the afferent neuron by closing K_V_7 channels on the calyx outer face (*COF*) (15). Cells within intact vestibular epithelia are densely packed with synaptic and non-synaptic extracellular spaces (*ECS*) that are continuous and of comparable width and K^+^ channels are present on the COF and at hemi-nodes (7, 16). This raises the possibility that dynamic changes in electrical potential and [K^+^] occur not only in the synaptic cleft, but throughout the ECS of the epithelium. We hypothesized that when one or more type I hair cells experience bundle deflection: 1) currents flow into ECS of the epithelium surrounding the COF to modulate local extracellular potentials and are in turn affected by them; 2) such extracellular potentials enhance AP recruitment in afferent calyces.

## Results

Based on our previous work (4), we developed a 3D finite element model to compare the *in vitro* condition where an isolated vestibular hair cell-calyx and the associated afferent are in a perilymph bath (VHCC, **Fig 1 A1**) with the semi-intact or *in vivo* situations where the synaptic cleft is continuous with an extensive network of ECS that lead to perilymph past basement membrane (VHCC-E, **Fig 1 A2**). The VHCC-E model thus represents a more realistic pathway for ion flow and allows for modulation of channels located on the COF. For a 1 μm excitatory hair bundle deflection, AP latency is reduced when simulated within the epithelium (**Fig 1B**). To distinguish whether electrical potential or K^+^ accumulation in the ECS were primarily responsible for the latency reduction, we added potassium chloride cotransporter (KCC) activity to the COF and supporting cell membranes - in addition to its location on the CIF in the VHCC model (4). Our epithelial model contained five adjacent hair cell-calyx complexes, as can occur in-vivo (especially in the striolar/central regions) (**Fig 2A**). When only the hair cell in the center is stimulated (**Figs. 1B and 2A**), a significant reduction in AP latency remained (**Fig 1B**) even when extracellular [K^+^] was reduced (**Fig 2A, top row**). Importantly, the extracellular electrical potential surrounding calyceal terminals was still significant (**Fig 2A, bottom row**). Extracellular potentials were greater in magnitude along the calyx membrane when within the epithelium (VHCC-E, **Fig. 2B**). We subsequently investigated simultaneous bundle deflection of all five hair cells with greater KCC activity in the epithelia. The type-I hair cells are labelled based on their distance to the cell in the center (C), as the 1^st^ (1N) and 2^nd^ nearest neighbors (2N) (inset, **Fig 3A1**). Excitatory deflection of the central hair bundle caused an AP to fire in the associated afferent and created small nonquantal EPSPs in neighboring afferents (**Fig 3A1, A2**). When all (C, 1N, 2N) hair bundles were deflected together, by either 1 μm or 0.4 μm, the AP latency in the “C” afferent was further reduced by 1.2 ms (**Fig 3B1**) or 5.2 ms (**Fig 3B2**) respectively.

**Fig 1.**
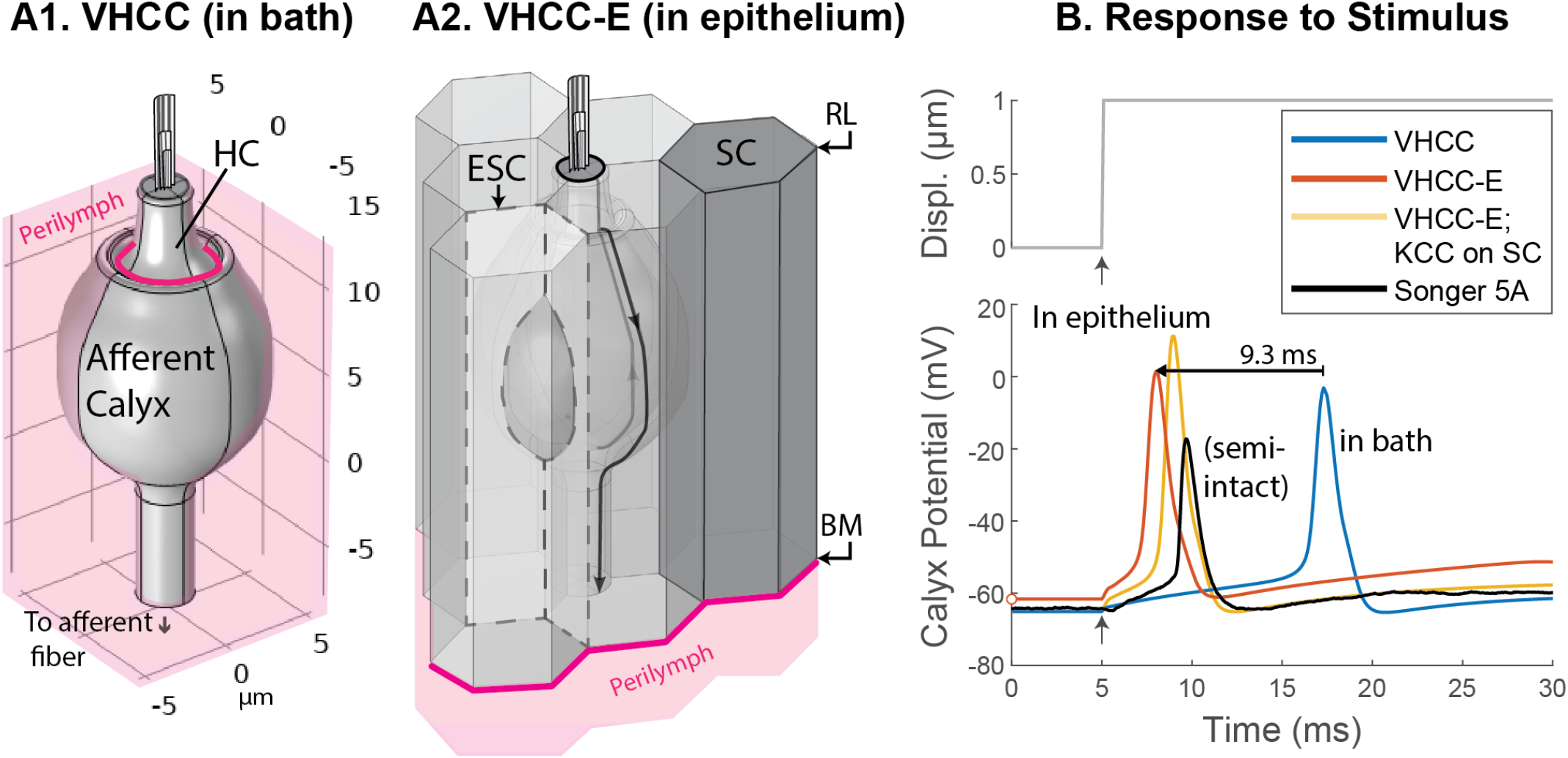
AP latency is reduced when cells are within the epithelium. A1. A large perilymph bath ([K^+^] = 5 mM and 0 mV) surrounds the entire hair cell and calyx and the perilymph boundary condition is applied at the lip of the calyx where the synaptic cleft would meet perilymph (pink curve); as in (Govindaraju et al. 2023), perilymph bathes the COF. **A2** In the epithelial model (VHCC-E) the perilymph boundary (pink curve) is at the basement membrane (BM) and electro-diffusion is calculated in both synaptic (between hair cell and calyx) and non-synaptic extracellular spaces between supporting cell (SC), the hair cell and the calyx. Within the epithelium, surfaces represent cell membranes and the adjoining extracellular space (ECS); examples are marked with dashed lines. **B** APs generated under VHCC and VHCC-E conditions are shown. The VHCC AP peak occurred 12.3 ms after stimulus onset (blue curve) and was greater than that seen experimentally (black curve, adapted from Songer and Eatock 2013, Figure 5A, 4.7 ms) in a semi-intact preparation. When dynamic changes in extracellular electrical potential and ion concentration are considered in the VHCC-E model, the AP peak (orange curve) occurred at 8 ms, a latency of 3 ms. When large amounts of a potassium chloride cotransporter (KCC) were situated on supporting cell surfaces (VHCC-E; KCC on SC – yellow curve) to minimize changes in extracellular [K^+^], the AP Peak occurred at 9 ms, a latency of 4 ms.

**Fig 2.**
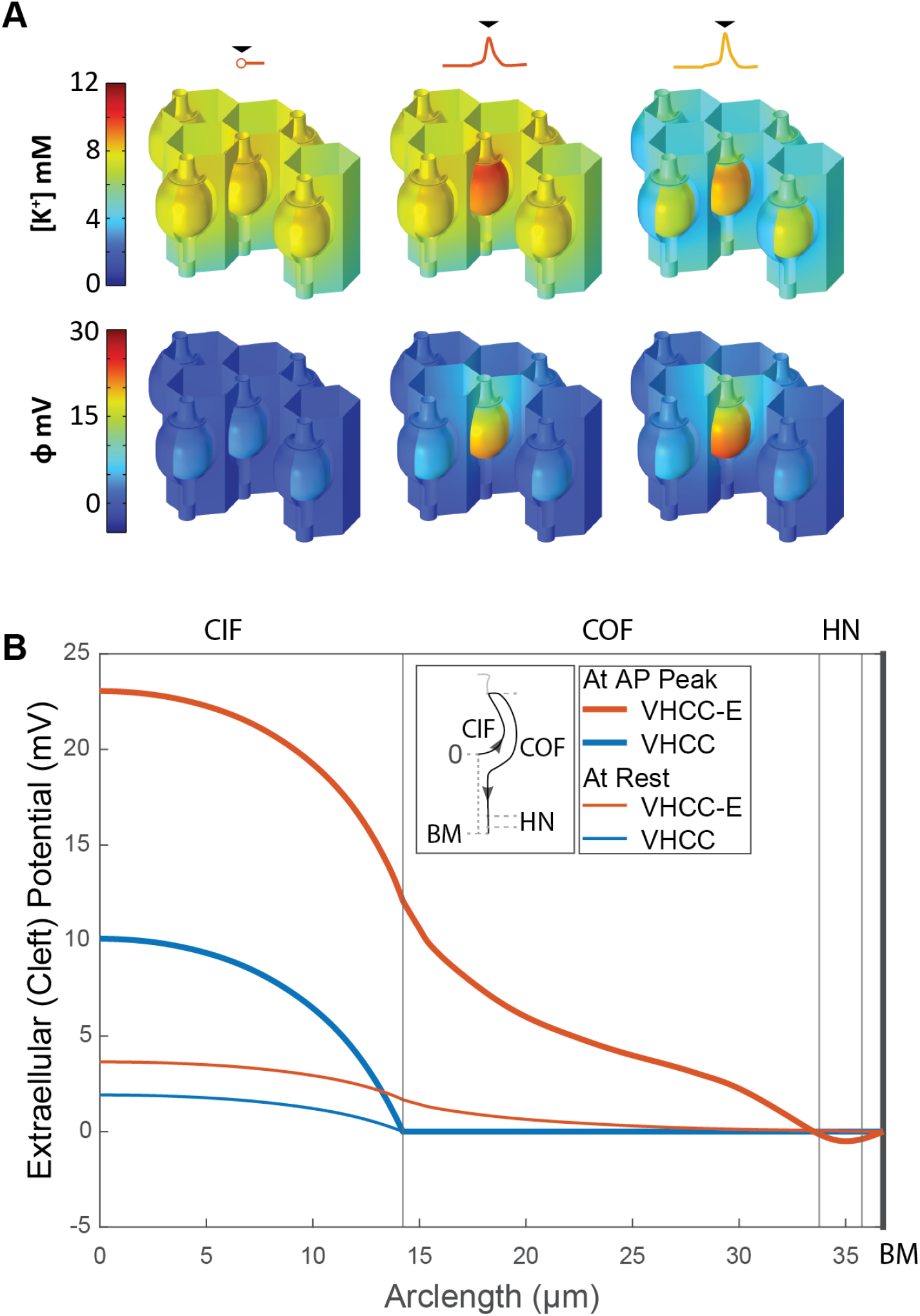
Extracellular electrical potential rises significantly during APs. **A**. The 3D spatial gradients in extracellular potassium ion concentration **[K**^**+**^] (top row) and electrical potential **ϕ** (bottom row) are shown at rest and at the AP peaks (black arrowheads) seen in Fig 1B where only the central hair cell is stimulated for the VHCC-E condition (2^nd^ column) and with KCC on SC (3^rd^ column). The outer surface (COF) of calyces in the foreground have been partially removed to show the synaptic cleft. **B**. The spatial profile of extracellular potential along the calyx from the base of the inner face (CIF) to the basement membrane (BM) is shown when only the central hair cell is stimulated. In the epithelial model (red lines, VHCC-E), the magnitude of the extracellular potential is increased both at rest (thin lines) and during the action potential (thick lines) with respect to a calyx in bath (blue lines, VHCC).

**Fig. 3.**
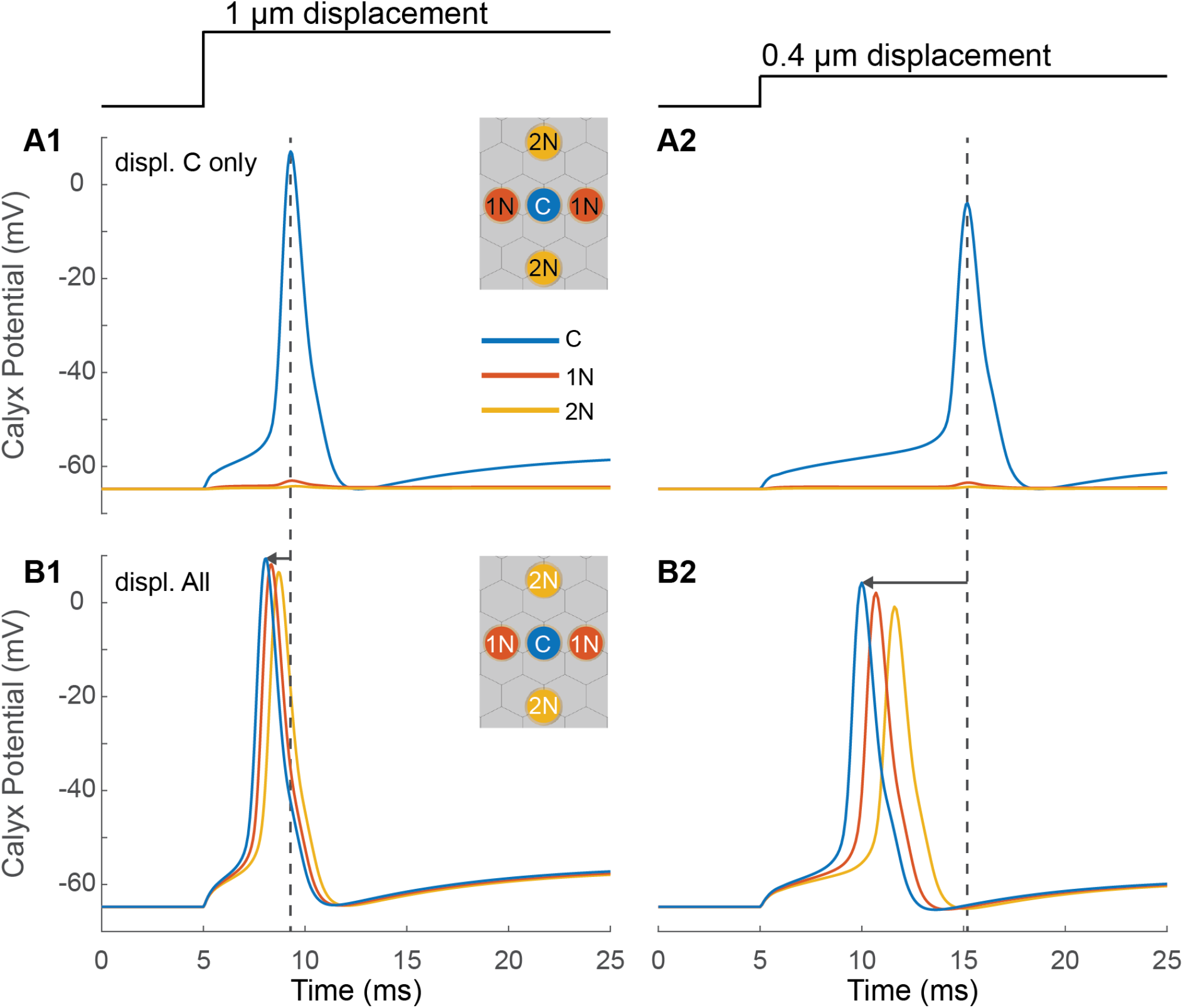
Collective hair bundle deflection reduces AP latency. When only the hair bundle of the center hair cell is displaced (**A1, A2**), the rise in extracellular potential surrounding the associated calyx, over the course of the AP (blue curve), caused small depolarizations in the 1st (1N) and 2nd (2N) nearest neighbors (yellow and orange curves). Inset: Top-down view of the arrangement of hair cells. When the hair bundles of all hair cells are displaced together, APs occur in all calyces (blue, yellow and orange curves) (**B1, B2**). The maximal reduction in AP latency is experienced by the calyx in the center. The improvement in AP latency due to concomitant depolarization of adjacent calyces is greatest for the smaller, 0.4 μm, displacement (1.2 ms vs 5.2 ms, arrows, **B1 vs B2**) although the larger displacement results in an AP that peaks at an earlier time (8 ms vs 10 ms, blue curve, **B1 vs B2**).

## Discussion

In the VHCC-E model, following excitatory bundle deflection, the afferent AP occurred with reduced latency (**Fig 1B**) and was closer to in-vitro experimental data (2) than when surrounded by a perilymph bath. The reduction in AP latency can be attributed, primarily, to greater changes in extracellular electrical potential during afferent depolarization (**Fig 2**) which alter the driving forces of currents across the CIF and COF. The voltage and ligand-gated K_V_7 channels (17) found on the CIF and COF are non-inactivating, depolarization-activated, and modulated by efferent activity. Increases in extracellular electrical potential reduce K^+^ efflux through these channels and notably reduce AP latency for smaller hair bundle displacements applied concomitantly to multiple adjacent hair bundles (**Fig 3**) – as would occur in-vivo during head motion. *In-vivo* extracellular recordings, in response to applied head motion stimuli, show shorter time to spike and encoding of higher frequency stimuli than seen *in-vitro* in recordings from an individual calyx terminal where only the hair bundle of the associated type-I cell is displaced (2, 13, 18). We posit that transient local potentials in the ECS act in concert with hair cell, calyceal and hemi-nodal currents to enhance transmission at the synapse and reduce the decay of the afferent graded potential. This appears to be a way to lower AP latency without significant disruption to subsequent repolarization and may explain differences observed between *in-vivo* and *in-vitro* recordings (in addition to other factors such as temperature, age, species and presence of endolymph). The model also rationalizes observations that calyx bearing afferents were sensitive to applied galvanic stimulation (extracellular current injection) (19). Our results suggest this sensitivity reflects a natural ability of calyceal afferents to create and interact with extracellular potentials, within the intact epithelium, to support faster AP generation during head motion.

## Methods

Simulations were performed in COMSOL 6.3. Governing equations were used as in our previous publication (4) with the electro-diffusion equation now applied to the entire extracellular space as opposed to just the synaptic cleft. Other modifications are detailed in the Supporting Information (SI).

## Acknowledgements

This study was supported by NIH-NIDCD (R21 DC022412 and R01 DC012347).

## Supporting Information

## Computational Methods and Modifications

### Overview

The model in this paper uses a finite-element-analysis simulation software (COMSOL), to solve equations for K^+^ and Na^+^ electro-diffusion in the synaptic cleft and extracellular space, Hodgkin-Huxley-style equations to represent voltage-dependent ion flow through ion channels, and the cable equation for electrical propagation. For each simulation, steady-state conditions are established and followed by a step deflection of one or more hair bundles. Outputs include the basal-to-apical gradients within the synaptic cleft and in the surrounding extracellular spaces of the epithelium for K^+^, Na^+^ and electrical potential, local membrane voltages across afferent neuron surfaces, including at the hemi-node and up to the first two full nodes of the distal branch of the bipolar vestibular afferent neuron. The distal branch connects the calyceal terminals to the neuronal cell body in the vestibular ganglion; the remainder of the distal branch, the cell body and the central branch projecting to the brain are not included in the model. Negative currents represent the flow of cations into the cell bearing the channel or transporter in its membrane. **The methods for the above mentioned processes and capabilities are presented in detail in supporting information of our previous publication in PNAS** (Govindaraju et al. 2023)**; below we describe notable changes made in the model used in this article**.

1. The use of the 3D geometry presented in the paper. A 3D representation of the hair cell-calyx geometry was created using the same measurements as before. Additionally, an array of hexagonal prisms representing supporting cells was created to surround the 3D hair-cell calyx geometries. The faces formed upon combining these structures were used to represent the narrow network of extracellular space that exists between cells in the epithelium. In simulations using the epithelia model (VHCC-E) five afferent calyx terminals with a hair cell each, were situated amidst a 120 μm x 126 μm patch of epithelia formed by an array of hexagonal prisms representing supporting cells. These prisms were 23.6 μm tall from base to apex, the diameter of the hexagonal cross-section was 12 μm. In Fig. 1B only cells adjacent to the calyx terminal in the center are depicted. In Figs. 2 and 3, only cell surfaces adjacent to the five calyx terminals are shown. Bundle displacement was applied to the center hair cell only or to all hair cells (illustrated in Fig 3).
2. Previously the Nernst-Planck electro-diffusion equation was applied only in the synaptic cleft to model changes in [K^+^] and [Na^+^]; we now additionally consider electro-diffusion in the non-synaptic extracellular spaces of the epithelium. For VHCC-E, the perilymph boundary condition (*ϕ* = 0 mV; [K^+^] = 5 mM; [Na^+^] = 140 mM) is moved to where the afferent fiber exits the epithelium, i.e. to the basement membrane.
3. The expression describing the dominant slow time constant (in ms) of the low voltage gated potassium conductance (gK,L) in our previous publication was replaced with an expression (S1) fitting time constant data from Rüsch and Eatock (1996). We found that these time constants improved the model’s ability to capture data seen in Lim et al. (2011) as described in Govindaraju (2023, Ph.D. Thesis). Additionally, we replaced the previous time constant for HCN with an expression (S2) from Meredith, Benke, and Rennie (2012).

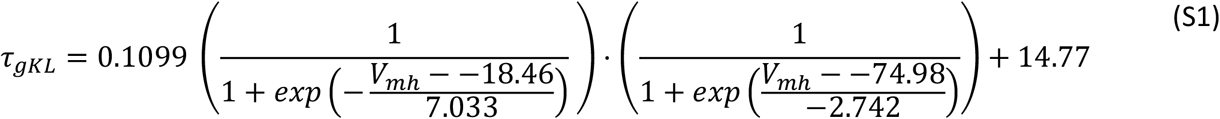

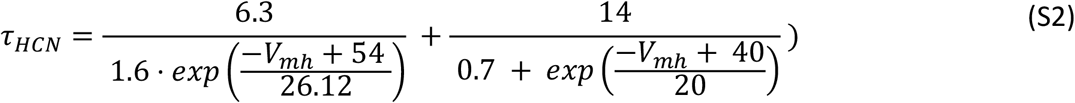
4. The hair cell was previously assumed to be equipotential based on its small overall size and its change in potential was described using an ordinary differential equation (S3) where the time dependence of

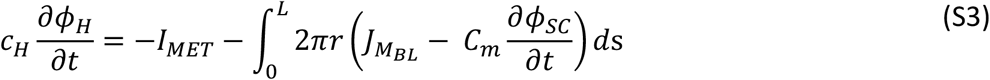

the hair cell electrical potential (*ϕ*_*H*_) is calculated as a function of hair cell capacitance (*C*_*H*_), mechanotransduction current (*I*_*MET*_), and current across the basolateral membrane 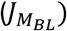. The integral represents the total current across the basolateral hair cell surface within the calyx. *L* is the length of the synaptic cleft from base to apex. *ds* is an infinitesimal arclength along the cleft. The above approach required the creation of an integration operator for each hair cell to calculate the corresponding basolateral current for use in the ODE and was not scalable when simulating

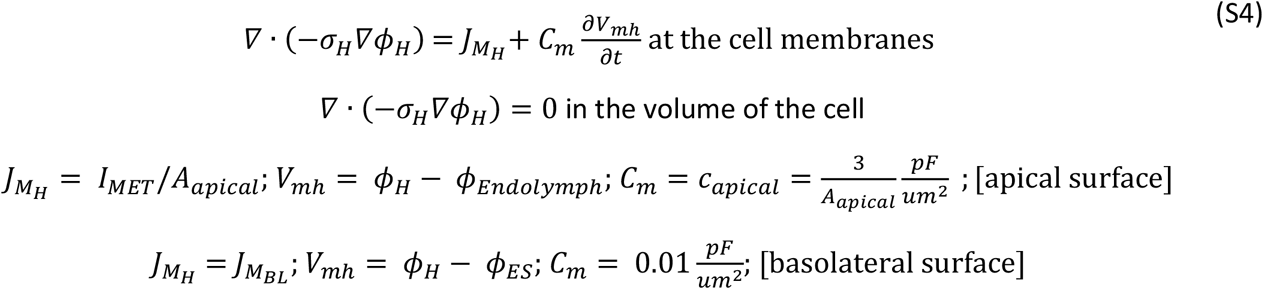

greater numbers of hair cells. Instead, we now use a partial differential equation (S4) that describes spatiotemporal changes in hair cell potential caused by temporal- and voltage-dependent changes in currents at the apical and basolateral cell membranes. *σ*_*H*_ is the conductivity of the hair cell, 1000 nS/μm (equivalent to 1 S/m) as previously used for the calyx and synaptic cleft space. *ϕ* _*Endolymph*_ is 5 mV, as used previously. 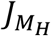 is the membrane current density. *A*_*apical*_ is 4.24 um^2^ for the hair cell geometry used in the model. At the apical surface the membrane voltage is the difference in potential between the hair cell (*ϕ*_*H*_) and endolymph (*ϕ* _*Endolymph*_); at the basolateral surfaces it is the difference in potential between the hair cell and the surrounding synaptic/non-synaptic extracellular space (*ϕ*_*ES*_). Given the diameter of the hair cell, differences in intracellular potential from apex to base were typically less than 0.5 mV.
5. Previously, we had distributed 30 nS/μm^2^ of Na_V_ conductance along an 8 μm unmyelinated section of fiber beneath the calyx and preceding the hemi-node. The hemi-node was 1 μm in length and had a Na_V_ conductance of 120 nS/μm^2^; the fiber diameter was 3 μm. In this model version, we retained an equivalent total conductance by using a value of 180 nS/ μm^2^ on a 2 μm long hemi-node for Figs. 1 and 2. Observations communicated by Dr. Anna Lysakowski showed that hemi-nodes of central/striolar calyces tended to occur within the basement membrane. For this reason, we situated the hemi-node at a distance of 5 μm from the base of the calyx inner face. When collectively stimulating all hair bundles, we observed some subthreshold oscillations following the AP when using a Na_V_ conductance density of 180 nS/um^2^. To clearly present the qualitative effects on first-spike timing during collective hair bundle displacement when in epithelia, we reduced Na_V_ conductance density at fiber hemi-nodes to 120 nS/um^2^ for the traces shown in Fig.3. This resulted in a 0.2 ms increase in time to peak for similar bundle displacement (Fig 3A1 vs Fig 1B VHCC-E with KCC on SC) and eliminated the subthreshold oscillations following stimulus. The spatial variations in sodium channel conductance density along the afferent fiber are not yet known although electrophysiological recordings demonstrate zonal and developmental differences in sodium currents in vestibular afferents (Meredith and Rennie 2018).

### Model Simulation and Solver Conditions

The surfaces of hair cells, calyx terminals and supporting cells were meshed, these surfaces were used to represent both the corresponding cell membranes and the adjoining extracellular space. The volumes within the hair cells and calyces were also meshed. In the epithelial model, there were 67753 tetrahedrons, 30774 triangles, 4750 edges, 866 vertices. The number and distribution of finite elements was chosen to minimize error in the solutions of the governing equations and to reduce computational time. To obtain the resting conditions, we first ran the model simulation with a stationary solver. We then used a time-dependent solver to determine the responses in the hair cell, synaptic cleft, calyx, and afferent fiber during a step displacement of the stereociliary bundle of one or more hair cells.

For step stimuli, the specified output time ranges were 0:0.1:30. The middle value indicates the increment between the time values (in ms) on either side. The time dependent solver in COMSOL utilizes adaptive time stepping which is also affected by an error tolerance setting within the software. A relative tolerance of 1E-4 was used. The solver has multiple settings which control aspects of adaptive time stepping and how steps are taken in relation to output times. Results are interpolated where solver time steps do not exactly coincide with specified output times. We used the ‘strict’ or ‘intermediate’ settings which ensure a step is taken at each specified output time. Enforcing smaller time steps for the full duration of the simulation did not significantly alter the computed results. The time-dependent solver was run using the backward difference formula (BDF) method.

The model simultaneously solves governing equations to simulate the electrical potential of the hair cell and the spatiotemporal evolution of electrical potential, potassium concentration, sodium concentration in the extracellular space (synaptic / non-synaptic), and potential in the afferent fiber.

**Model Repository**: https://github.com/acgsci/VestibularEpitheliaModel

The simulation files used to generate the presented results are available at the link above.

## Notes

https://github.com/acgsci/VestibularEpitheliaModel

### Competing Interest Statement

The authors have declared no competing interest.

